# A Sex-Specific Switch in Platelet Receptor Signaling Following Myocardial Infarction

**DOI:** 10.1101/580415

**Authors:** Beom Soo Kim, David A. Auerbach, Hamza Sadhra, Frederick S. Ling, Sandra Toth, Amy Mohan, Sara Tura, Ilan Goldenberg, David Q. Rich, Scott J. Cameron

## Abstract

**BACKGROUND:** A Sex-specific, personalized approach to anti-platelet therapy may be important in patients with myocardial infarction (MI).

**OBJECTIVES:** Our goal was to determine whether platelets activate differently in healthy men and women compared to following MI.

**METHODS:** Blood was obtained from healthy subjects or patients presenting acutely with ST-segment Elevation Myocardial Infarction (STEMI) and non-ST Segment Elevation Myocardial Infarction (NSTEMI). Platelet function through surface receptor activation was examined in healthy subjects, in patients with MI, and in age- and strain-matched mice before and after MI. Multivariate regression analyses revealed clinical variables associated with platelet receptor sensitivity at the time of MI.

**RESULTS:** Platelets from healthy women are dose-dependently more active compared to men, particularly through the platelet thromboxane signaling pathway (7.8-fold increase in women vs. 3.0-fold in men, P=0.02). At the time of MI, platelet activation through surface protease-activated receptor 1 (PAR1) was less in women than men (3.5-fold vs. 8.5-fold, respectively, P=0.0001). Multivariate regression analyses revealed male sex (P=0.04) and NSTEMI (P=0.003) as independent predictors of enhanced platelet PAR1 signaling at the time of MI. Similar to humans, healthy female mice showed preferential thrombin-mediated platelet activation compared to male mice (8.7-fold vs. 4.8-fold, respectively; P<0.001). In the immediate post-MI environment, male mice showed preferential thrombin-mediated platelet activation compared to female mice (12.4-fold vs. 5.5-fold, respectively; P<0.001).

**CONCLUSIONS:** These results outline a previously unrecognized sex-dependent platelet phenotype where inhibition of thrombin signaling in the peri-MI environment—particularly in males—may be an important consideration.

**CONDENSED ABSTRACT:** Preclinical studies evaluating anti-platelet drugs are generally conducted in platelets isolated from healthy individuals. Growing evidence suggests changes in platelet signaling properties in certain disease conditions compared to healthy platelets may alter the response to anti-platelet medications. This investigation revealed that platelets from men and women who are healthy and following MI signal differently, particularly through thromboxane and PAR1 receptors. This effect was especially noted in patients with NSTEMI compared to STEMI. These observations raise the possibility of considering a sex-specific anti-platelet regimen for males and females in atheroembolic vascular diseases such as NSTEMI.

## INTRODUCTION

Anti-platelet therapy for patients following a diagnosis of MI consists of aspirin and a P2Y_12_ receptor antagonists irrespective of whether the diagnosis is STEMI or NSTEMI. Recent data conducted in humans and in relevant murine models of MI and peripheral arterial disease (PAD) suggests the platelet phenotype, platelet receptor signaling, and responsiveness to anti-platelet medications are quite different from healthy platelets in which anti-platelet agents are first characterized (1–6).

Patients with established vascular disease have a heightened risk of recurrent atherothrombotic events, often manifesting as (NSTEMI) (7). Observational studies indicate long-term mortality for patients with NSTEMI remains greater than STEMI irrespective of appropriate cardiac catheterization and revascularization utilization (8,9). We previously reported in a small patient cohort that platelets from patients with NSTEMI show differences in post-receptor signal transduction pathways including the platelet PAR1 pathway compared to platelets from patients with STEMI or from healthy individuals (1).

Cardiovascular physiology and pharmacological responses in men and women are not the same (10). Myocardial function and the ability to withstand ischemic insult may also differ in women compared to men (11–14). Women who present with acute and chronic cardiovascular diseases remain under-studied and, therefore, are likely under-treated (15). The clinical presentation, management, and response to therapies for MI may also be different in women compared to men (16,17). Women with NSTEMI are less likely to be treated with an early invasive strategy than men though, curiously, this did not contribute to increased mortality in women (18).

Anti-platelet therapy prescribed following acute MI or as maintenance therapy in patients with coronary artery disease (CAD) does not consider whether the patient is male or female. A recent meta-analysis suggested that low dose aspirin may be less efficacious in men compared to women based on weight and height being associated with adverse vascular events or bleeding diatheses (19). We rather hypothesized that sex-specific differences in platelet signaling in health and at the time of MI may account for such population-based observations. To test this hypothesis, we evaluated platelet signaling following agonist stimulation in healthy women and men, comparing those responses to platelet signaling in women and men presenting with acute MI. We then interrogated our data to determine independent clinical variables predicting platelet responses though specific surface receptors for which prescription medications exist.

## METHODS

### HUMAN STUDIES

Healthy volunteer subjects aged 19-89 were recruited by answering a posted document, and all were free from anti-platelet agent use or vasoactive substances. The post-MI patient cohort included STEMI and NSTEMI patients. Delayed consent was granted for patients presenting with STEMI (within 24 hours of presentation) to avoid interfering with the door-to-balloon time. This also provided platelets obtained before they were affected by P2Y_12_ receptor antagonist load, and before angiography or coronary instrumentation. The sample was discarded if the patient declined to enroll and sign the consent form. For patients with NSTEMI, subjects were already in the emergency department or hospital for 1-10 hours and identified by elevated plasma cardiac troponin (plasma value greater than the 99^th^ percentile of a healthy population with a 10% assay coefficient of variation CV) and symptoms consistent with cardiac chest pain. Each patient was treated with 325 mg aspirin in the emergency department or the ambulance at least 30 minutes prior to venous blood draw. We enrolled 143 individuals: 26 healthy volunteers, and 117 patients with acute MI. Patients were de-identified and basic patient demographics and clinical characteristics were collected. This study was approved by the University of Rochester Research Subjects Review Board (RRRB) protocols 61784 (patients with MI) and 28659 (healthy subjects).

### PLATELET FUNCTION

We used flow cytometry with measurement of platelet surface P-selectin to assess platelet function in a dose-dependent manner, as we have described previously (1). Platelet surface P-selectin is a marker of platelet alpha granule exocytosis in response to pharmacologic doses of platelet receptor agonists: ADP (P2Y_12_ receptor), TRAP-6 (PAR1), and U46619 (thromboxane receptor). For the purposes of interrogating platelet activity, this technique is a reliable marker of platelet activation and shows similar data to light transmission aggregometry (20). Blood was collected by a trained medical professional into citrate plasma tubes. Analysis occurred within 60 minutes of blood collection. Platelet rich plasma (PRP) was isolated, with each concentration of agonist stimulation performed in quadruplicate. Following 15 minutes of agonist stimulation, 1 μL of labeled CD62P (P-selectin) antibody was incubated in the dark for 30 minutes. Samples were fixed in 2% formalin, then platelet surface P-selectin was quantified on an Accuri Flow Cytometer (BD Biosciences). Data was then processed through FloJo (Ashland, Oregon).

### PLATELET SPREADING

Glass slides were pre-coated the prior day with fresh human fibrinogen at a final concentration of 2.5 mg/mL in PBS. The fibrinogen-coated slides were washed three times with PBS and then incubated for 1 hour at 37°C with PRP from healthy subjects. Platelets were then fixed in 4% formaldehyde in 0.25% Triton X-100 for 20 minutes and then incubated with 0.3 μM rhodamine phalloidin for 30 minutes. The glass slides were then gently washed three times with PBS, and then imaged with a confocal microscope. The investigator was blinded to the identity of the platelets until the analysis was complete and decoding occurred. Random fields of platelets were imaged, and the average platelet area was calculated using ImageJ software (NIH).

### REAGENTS

TRAP-6 (Cayman Chemicals # 3497), 2-methyl-ADP (Tocris, Bristol, UK # 475193-31-8), U46619 (Cayman Chemical # 56985-40-1), Thrombin (Sigma # 9002-04-4). CD62P-PE antibody Clone AK4 # 12-0628-62 (Thermo Fisher Waltham, MA # F7496), CD62P-FITC (BD Pharmigen # Bdb553744) and FITC-Fibrinogen antibody (BD Pharimigen # F7496), P2Y_12_-FITC (Alamone #Apr-020-f), PAR1-FITC (LSBio #LS-C395810), FITC-thromboxane (Cayman Chemical #10012559).

### EXPERIMENTAL ANIMALS

Mouse colony: All animal protocols were approved by the University Committee on Animal Resources (UCAR). Eight-week-old male and female wild-type C57/BL6 mice were used in this study. The left anterior descending coronary artery (LAD) was surgically ligated as we previously described to create a model of chronic ischemia which chronically activates platelets over several days(2). Retro-orbital blood collected into heparinized Tyrodes solution as described by us previously was used to isolate mouse PRP(2). Methylene blue was injected and LV perfusion and ischemic area of risk confirmed the integrity of the model as we reported previously (2).

### STATISTICAL ANALYSIS

Dichotomous clinical variables are presented as frequencies. Continuous clinical variables are presented as mean with standard error of the mean (SEM) unless otherwise stated. Normalcy of all data were firstly evaluated by the Shapiro–Wilk test. For Gaussian-distributed data between 2 comparative groups, the t-test was used to assess for a difference between groups, and for non-Gaussian-distributed data, the Mann-Whitney *U* test was used. For Gaussian-distributed data in 3 or more groups, 1-way ANOVA then the Bonferroni multiple comparisons test was used, otherwise the Kruskal–Wallis test followed by Dunn post-test was used. Multivariate regression analyses were conducted in the post-MI cohort to identify clinical variables independently associated with platelet activation. Three separate models were performed to test for male/female and NSTEMI/STEMI specific differences in platelet reactivity through PAR1, the thromboxane receptor, and the P2Y_12_ receptor. The model included the following variables: sex, STEMI or NSTEMI (MI), age (rounded to nearest decade), hyperlipidemia (cholesterol), chronic kidney disease (renal function), chronic venous insufficiency (venous disease), hypertension (blood pressure), presently on platelet inhibitors (anti-platelet therapy), and peripheral artery disease (arterial disease.) Results include the parameter estimate ± 95% confidence interval (C.I.). Significance was accepted as a P value <0.05. Analyses were conducted using SAS version 9.4 (SAS Institute, North Carolina). Graphical data and simple group comparisons were illustrated using GraphPad Prism 7 (GraphPad Software, Inc., La Jolla, CA).

## RESULTS

### Sex differences in platelet signaling in healthy humans

Platelet signaling in healthy individuals was interrogated by dose-response curves using platelet surface agonists for which orally-available antagonists are available. Platelet PAR1 receptor signaling was assessed by TRAP-6, the thromboxane receptor by U46619, and the P2Y_12_ receptor by ADP. Platelets from women generally displayed more agonist responsiveness than men through the platelet receptors studied. The platelet thromboxane signaling pathway, which is indirectly inhibited by aspirin, was strikingly more active in platelets from healthy women compared to men (Figure 1). Spreading of human platelets on an extracellular fibrinogen matrix, which is suggestive of glycoprotein IIb/IIIa (GPIIb/IIIa) activativity, was enhanced in women compared to men (Figure 2).

**Figure 1.**
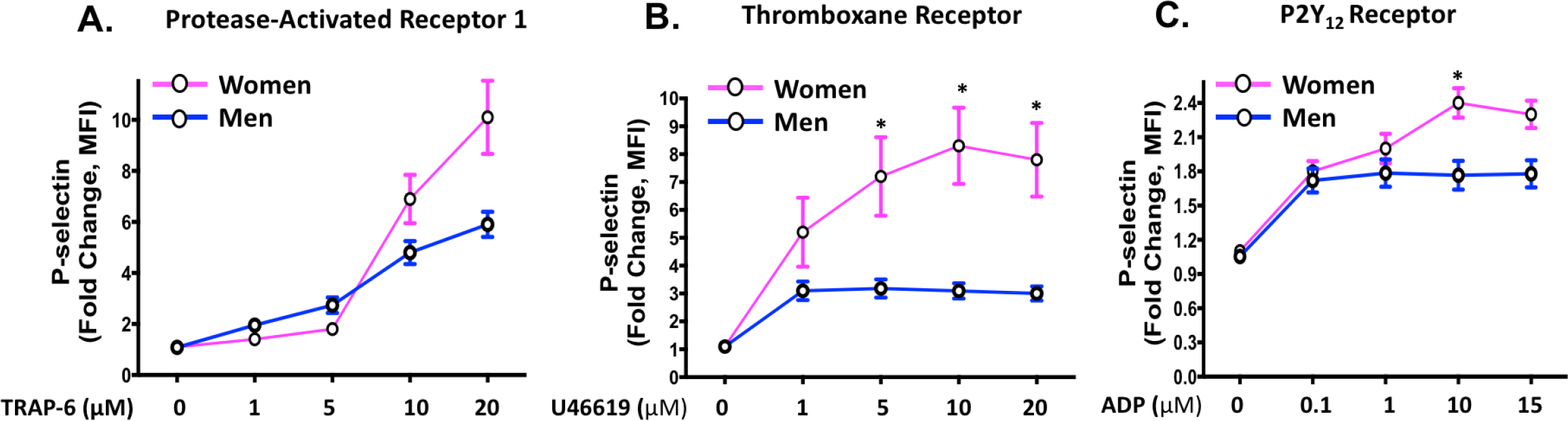
Agonist-stimulated platelet activation in healthy subjects: Platelets from healthy women and men were isolated and stimulated *ex vivo* with agonists for **(A)** PARl (TRAP-6), **(B)** the thromboxane receptor (U46619), and **(C)** the P2Y_12_receptor (ADP). Platelet activation was assessed by FACS for alpha granule secretion (surface p-select in) as mean fold increase from baseline ± SEM. * P <0.05 between groups at the indicated time point by the Kruskal-Wallis test followed by Dunn’s post test correction.

**Figure 2.**
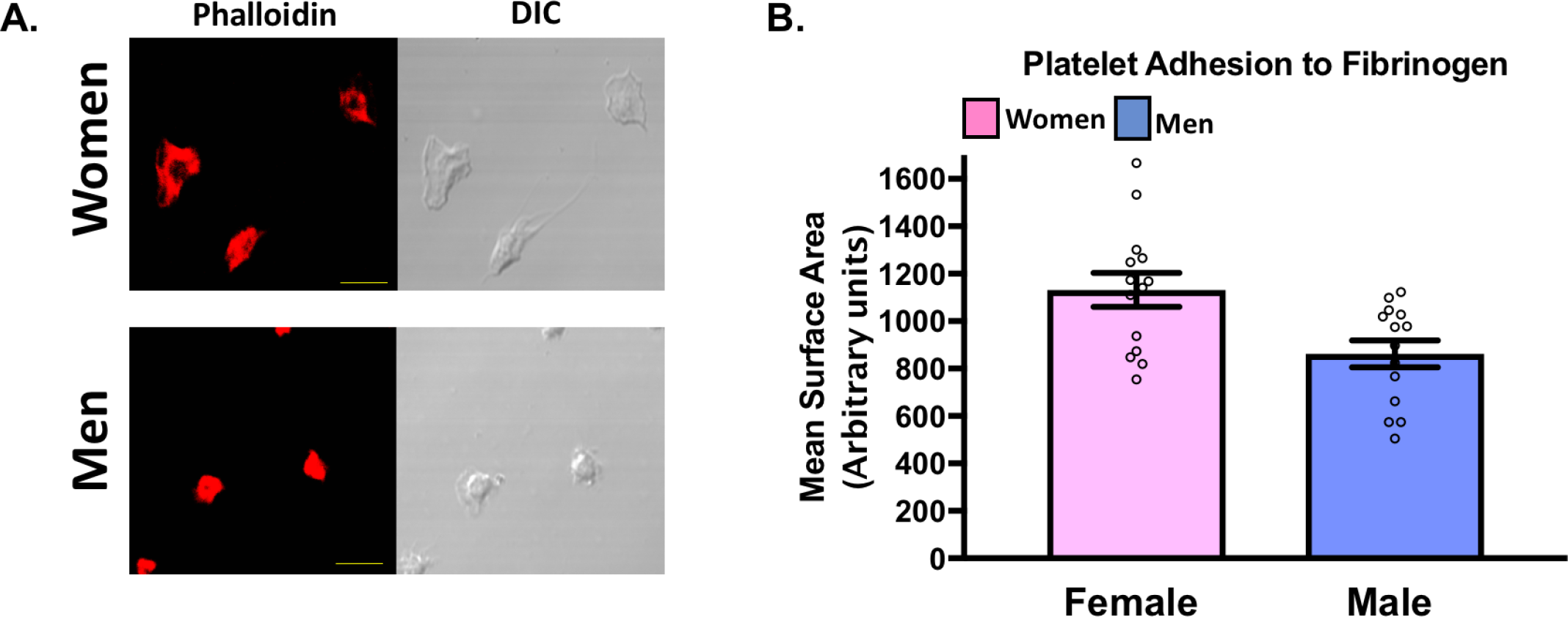
Spontaneous platelet activation in healthy subjects: **A.** Male and female platelets were assessed for adhesion to fibrinogen after 30 mins at 37 °C by confocal microscopy. **B.** Data are represented as mean platelet surface area± SEM. (n=3 in each group, 4-5 random fields per subject, 10-20 platelets per field). P=0.007 between groups, t-test. Phalloidin-PE stains actin cytoskeleton red. Yellow scale bar= SµM.

### Sex differences in platelet signaling post-MI in humans

Using a single platelet receptor agonist concentration which best evoked platelet responses in healthy platelets, we examined platelet activation in women and men presenting acutely with STEMI and NSTEMI prior to receiving a loading dose of a P2Y_12_ receptor antagonist and prior to coronary angiography. Table 1 shows the baseline clinical characteristics of the study population. Except for HDL, the clinical and demographic variables were similar in NSTEMI vs. STEMI and Male vs. Female participant groups. Contrary to our observation in healthy platelets, platelets in the immediate post-MI environment were more reactive through platelet PAR1 by 2.7-fold in men compared to women (P=0.001), and 1.6-fold more reactive for men compared to women through the platelet P2Y_12_ receptor (P=0.04). There was no difference in platelet activation between men and women through the platelet thromboxane receptor following MI (Figure. 3).

**Table 1.** Demographics of patients presenting with MI: Demographic characteristics of the post-Ml study population. When stratifying for sex and NSTEMI/STEMI, except for HDL, the study populations were similar. There were 60 NSTEMI and 57 STEMI patie nts, and males comprised a similar proportion of each of these cohorts {30-33%.) Data are shown as mean ± SEM. Level of significance is noted. HDL=high densit y lipoprotein. LDL=low density lipoprotein. LVEF=left ventricular eject io n fraction. All data presented reflect variables at the time of patient evaluation in the emergency department.

| Clinical Characteristics | NSTEMI | STEMI | p-value |  | Male | Female | p-value |
| --- | --- | --- | --- | --- | --- | --- | --- |
| Number of Patients (n) | 60 | 57 | ----- |  | 80 | 37 | ----- |
| Female, n (%) | 20(33) | 17(30) | 0.731 |  | 0 | 37(100) | <0.001 |
| Age at Admission (years) | 64.87±12.29 | 61.68±13.53 | 0.167 |  | 62.71±12.04 | 64.89±14.69 | 0.408 |
| Caucasian, n (%) | 54(90) | 43(77) | 0.055 |  | 67(84) | 31(84) | 0.996 |
| A1C (%) | 6.39±1.52 | 6.39±1.95 | 0.452 |  | 6.18±1.44 | 6.87±2.21 | 0.062 |
| Hyperlipidemia, n (%) | 45(75) | 43(77) | 0.822 |  | 63(80) | 25(68) | 0.153 |
| Hypertension, n (%) | 49(82) | 46(82) | 0.947 |  | 64(81) | 31(84) | 0.718 |
| History of Smoking, n (%) | 32(53) | 37(66) | 0.163 |  | 50(63) | 19(51) | 0.222 |
| Chronic Kidney Disease (%) | 45(75) | 50(88) | 0.078 |  | 64(80) | 31(84) | 0.626 |
| Platelet inhibitor, n (%) | 28(47) | 27(48) | 0.868 |  | 37(47) | 18(49) | 0.855 |
| Platelet count (x10<sup>3</sup>) | 208.28±50.48 | 239.89±93.38 | 0.133 |  | 218.96±72.44 | 232.76±81.83 | 0.433 |
| LVEF | 49.86±9.49 | 47.43±11.19 | 0.294 |  | 49.51±9.72 | 46.91±11.58 | 0.323 |
| Peak Troponin (ng/mL) | 0.91±0.34 | 1.89±0.37 | 0.608 |  | 1.41±0.45 | 1.41±0.26 | 0.998 |
| LDL (mg/dL) | 101.69±37.62 | 107.18±39.09 | 0.445 |  | 103.50±39.33 | 106.19±36.21 | 0.771 |
| HDL (mg/dL) | 45.81±16.40 | 53.10±18.25 | 0.022 |  | 46.17±15.42 | 56.52±20.32 | 0.017 |
| Statins, n (%) | 23(38) | 27(49) | 0.245 |  | 33(42) | 17(46) | 0.713 |
| Stroke, n (%) | 9(15) | 6(11) | 0.492 |  | 9(11) | 6(16) | 0.555 |
| Peripheral artery disease, n (%) | 5(8) | 3(6) | 0.72 |  | 3(4) | 5(14) | 0.107 |

**Figure 3:**
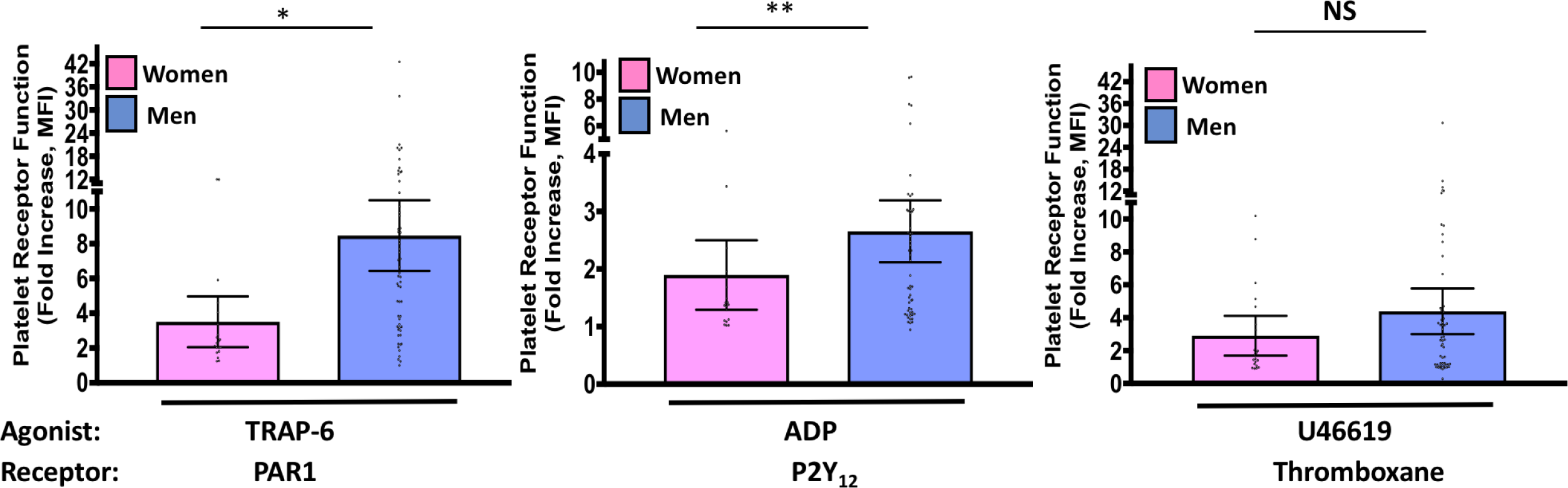
Platelet receptor function in patients with M.I. Blood was drawn from women and men as they were diagnosed with STEMI or NSTEMI, prior to coronary angiography, and prior to loading with a P2Y_12_ receptor antagonist. Platelet rich plasma was isolated and platelets were stimulated for 15 mins with an agonist for: PARl (TRAP-6, 10 µM), the P2Y_12_ receptor (ADP, 10 µM), or the thromboxane receptor {U46619, 10 µM). Platelets were labeled with a tagged antibody for P­ selectin, and then analyzed by flow cytometry. Platelet function is represented as median fold change in surface P-selectin as mean fluorescence intensity (MFI) from baseline± 95% C. I. Differences between women and men for each agonist was assessed by the M ann-Whitney U test. *P=0.0001. **P=0.0473. NS=Not significant.

A side-by-side comparison of platelets from healthy individuals compared to those at the time of MI revealed platelet PAR1 signaling decreased in women (P=0.031) and increased in men at the time of MI (P=0.0083, Figure S1). Similarly, platelets from women at the time of MI were less active through the P2Y_12_ receptor compared to platelets from healthy women (P=0.028). Platelets were slightly more active in men through the P2Y_12_ receptor at the time of MI compared to healthy men (p=0.055).

Multivariate regression analyses revealed that STEMI compared to NSTEMI was negatively associated with platelet thromboxane receptor function (PE −2.31, 95% CI −4.54 to – 0.07, p=0.04.). The presence of dyslipidemia (PE −5.18, 95% CI −9.29 to −1.06, P=0.01), female sex (PE −3.35, 95% CI −6.52 to −0.18, P=0.04), and STEMI (PE −4.71, 95% CI −7.7 to −1.7, P=0.003) were negatively associated with platelet activation through PAR1 (Figure 4). The observation that NSTEMI is more prominent in men and the strongest clinical predictor of platelet PAR1 signaling aligns with our previous observation in patients with NSTEMI (1), and contrasts what we observed in healthy individuals where women showed enhanced platelet PAR1 signaling (Figure 1). None of the clinical variables—including patient sex—were independently associated with platelet P2Y_12_ receptor function.

**Figure 4.**
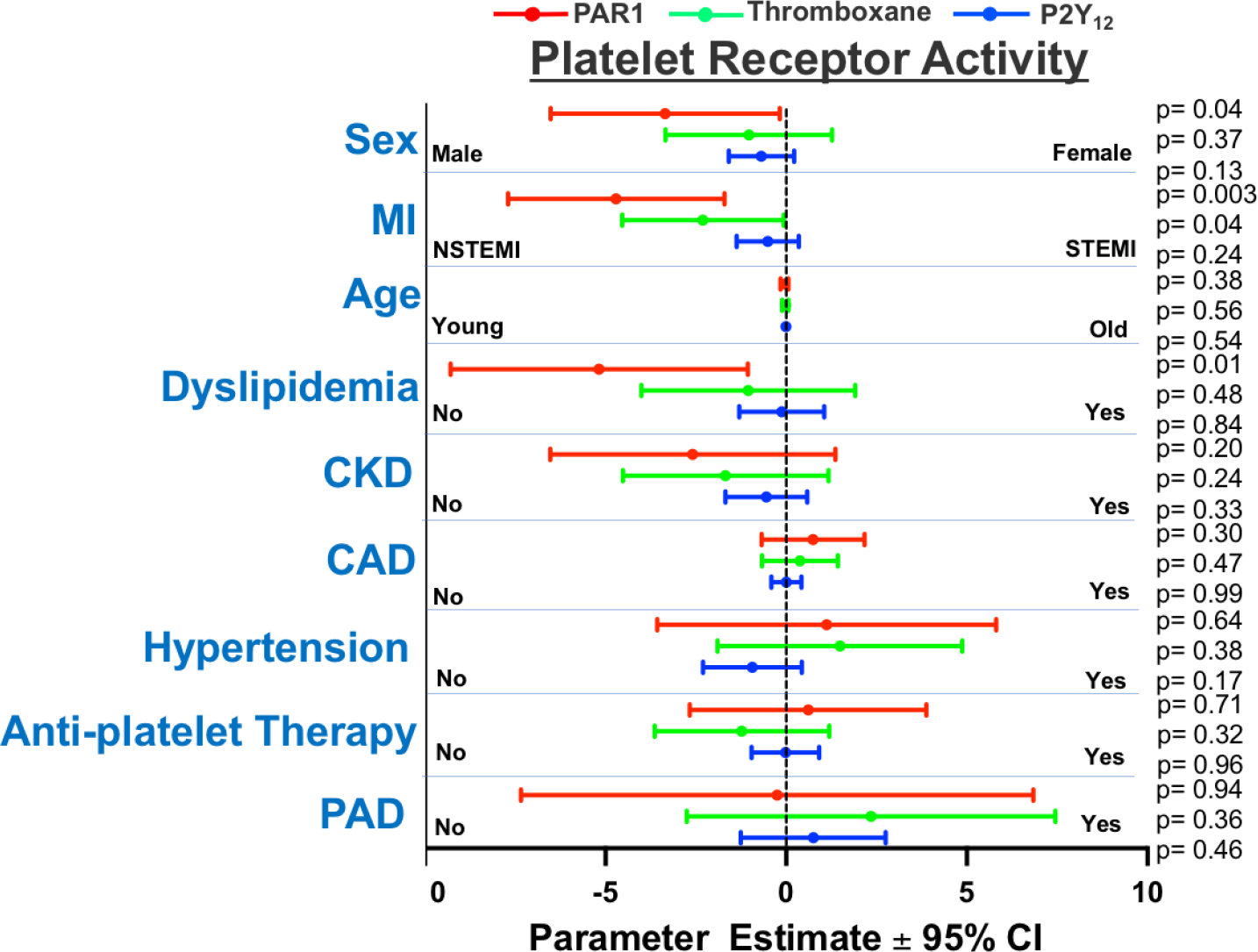
Independent Predictors of Platelet Receptor Activity after Ml: Forest plot illustrating the multivariate-adjusted mean hazard rat ios ± 95% C.I. of mult ivariat e-adjuste d regression analyses of for platelet function in response to surface receptors agonists at 10 µM. Red=PARl, Green=Th rombo xa ne, Bl ue=P2Y_12_. Level of significance is noted. There were 60 NSTEMI and 57 STEMI pat ie nts. CKD=c hronic kidney disease. CAD=ex isti ng coronary artery disease. PAD= peripheral artery disease.

**Figure 5A.**
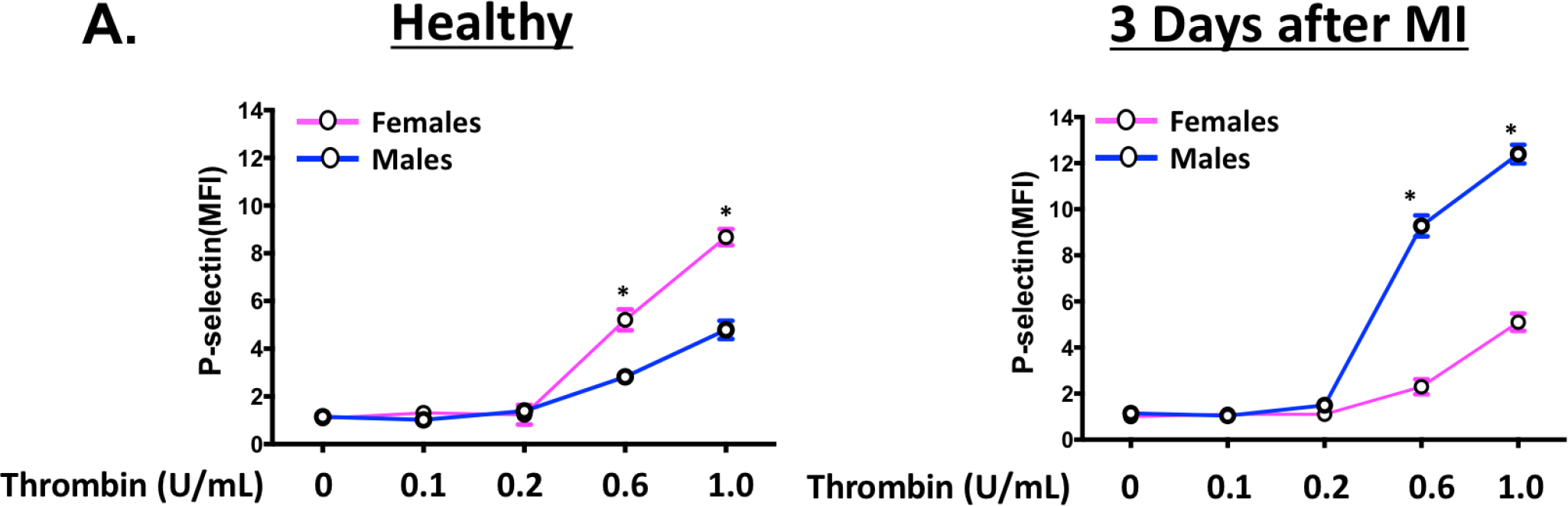
Thrombin-mediated platelet activation in mice before and following MI: Male and female C57/BL6 mouse platelets were isolated before and then 3 days after Ml and platelets were stimulated *ex vivo* with thrombin. Platelet activation was assessed by FACS (surface p-selectin). Data are represented as mean fold increase over baseline (O) ± SEM for n=S mice, each performed in quadruplicate. * P <0.001 between groups, assessed one-way ANOVA followed by Bonferroni’s multiple comparisons test.

**Figure 5B.**
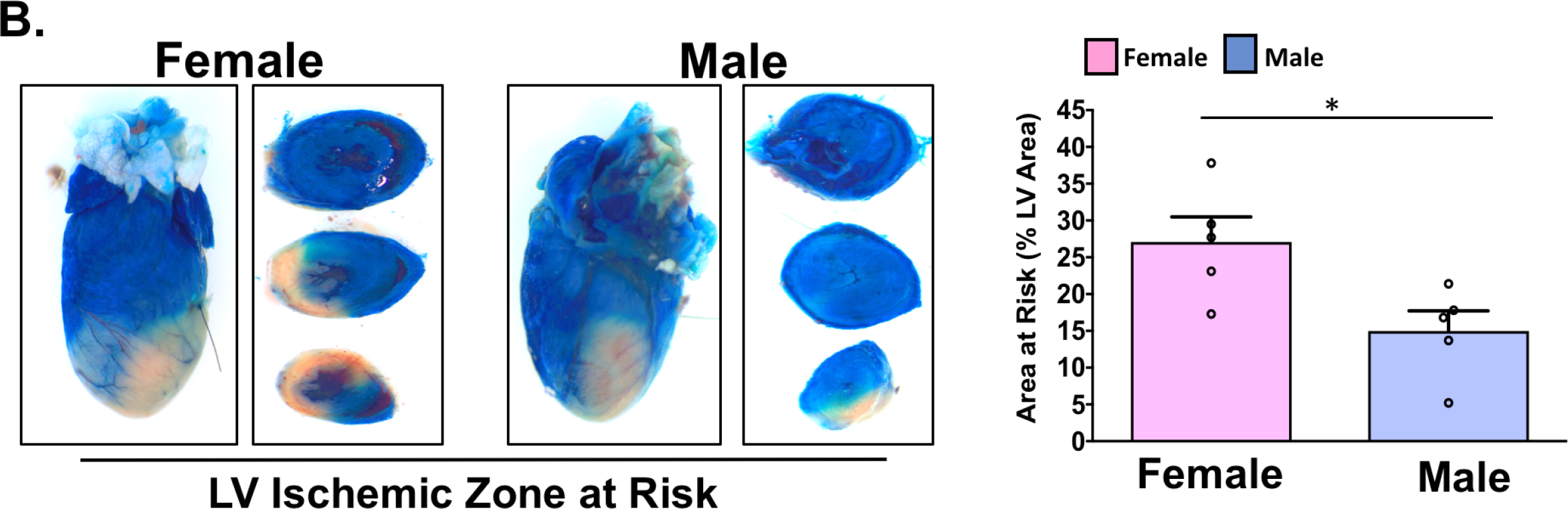
lschemic LV in mice after Ml: Male and female C57/BL6 mice underwent sham surgery or ligation of the LAD, followed immediately by retro-orbital infusion of 2% methylene blue to indicate areas of LV perfusion. Perfused tissue is blue and ischemic t issue pink. LV region at risk quantified as% LV area (mean% of LV area± SEM). Representative images are shown for male and female mice. LAD=left anterior descending coronary artery. LV=left ventricle. Differences between groups was assessed by t-test. *P=0.024.

### Sex differences in platelet surface receptor expression in humans

Using flow cytometry, we assessed whether differences in platelet surface PAR1, thromboxane, and P2Y_12_ receptor expression were responsible for the observed changes in platelet activation in women and men in health and following NSTEMI. Mild differences in platelet PAR1 receptor expression were noted with increased expression in stimulated healthy men(P=0.02) but not healthy women (P=0.08) platelets, and in both stimulated male NSTEMI platelets (P=0.023) and in stimulated female NSTEMI platelets (P=0.019). Platelet PAR1 expression in unstimulated healthy platelets compared to unstimulated NSTEMI platelet was similar in men and mildly increased in women (P=0.03) (Figure S2). Alterations in platelet surface receptor expression, therefore, is less likely to explain the profound increase in platelet PAR1 signaling observed in men presenting with NSTEMI (Figures 3-4).

Platelet surface thromboxane receptor expression was only greater in unstimulated NSTEMI platelets isolated from men compared to unstimulated healthy platelets (P=0.01), and for stimulated healthy platelets compared to stimulated NSTEMI platelets in men (P=0.04) (Figure S3). Only stimulated male NSTEMI platelets showed greater P2Y_12_ surface expression (105 vs. 61 MFI, P=0.045) (Figure S4).

### Sex differences in platelet signaling in healthy mice and in mice following MI

The mature platelet has a surprisingly dynamic protein expression profile which may be responsible for changes in platelet phenotype. Based on changes in agonist sensitivity, the response to anti-platelet agents, and platelet proteomic profiles, we concluded that the mature platelet phenotype can change *in vivo* and *in vitro* in humans and in mouse models of hypoxic and ischemic disease (21). Given our present observation that platelet thrombin signaling differs in men and women in health and following MI, we employed an established murine MI model previously found to have ongoing thrombin-mediated platelet activation in the post MI environment (2). Using each mouse as its own control, we compared platelet activation in male and female mice in the immediate pre-MI (healthy) environment to the post-MI (ischemic) environment. Consistent with our observation in human platelets, healthy female mouse platelets compared to male platelets preferentially signal through thrombin receptors (MFI 8.68-fold over baseline ± 0.34 vs. 4.79-fold over baseline ± 0.38, P=0.0002) and, 3 days after MI, were less reactive than males through thrombin receptors (MFI 12.4-fold over baseline ± 0.4 vs. 5.1-fold over baseline ± 0.38, P=0.0002) (Figure 5A). This observation is particularly striking given the area at risk in the ischemic left ventricle was slightly greater in females at the time of MI compared to males. This suggests an intrinsic difference in platelet signaling for males and females in the disease rather than the severity of the disease model (Figure 5B).

## DISCUSSION

To our knowledge, the present study is the first to directly compare platelet signaling in women with men in healthy conditions and in the post-MI environment. We purposefully investigated only platelet signaling pathways for which anti-platelet and anti-thrombotic agents exist to maximize the translational impact of the study. Comparing platelet signaling in health to the immediate post-MI environment, a reciprocal switch in platelet activation through protease-activated receptors occurs, with attenuated signaling in women and increased signaling in men during MI. This was further corroborated by reproducing this sex-specific observation in mice, suggesting a conserved rather than a species-specific physiologic phenomenon.

Our sex-specific observations in platelet signaling following MI was most pronounced in NSTEMI compared to STEMI, validating previous reports that platelets in each condition may be fundamentally different (1,22,23). By multivariate regression analysis, it is potentially revealing that NSTEMI independently predicted both platelet PAR1 and thromboxane, but not P2Y_12_ receptor signaling. This suggests that blocking platelet thromboxane production with aspirin and blocking the platelet PAR1 receptor directly with an antagonist such as vorapaxar, or indirectly with an anticoagulant, may be most beneficial in NSTEMI as implied in the original COMPASS study (24).

Women have historically been under-studied in scientific investigations. We confirmed the observation of Becker *et al.* that thromboxane-mediated platelet activation in health is more significant for women than men (25). Well-designed population-based studies failed to show that aspirin confers significant protection from adverse cardiovascular events in men when used for primary preventive purposes (26,27). When used for primary prevention in females, aspirin does, however, confer some protection against thrombotic stroke (28). A recent meta-analysis of five studies by Rothwell *et al*. suggested low dose aspirin may lack a protective effect in the majority of men due to possible under-dosing (19). While body mass and height tend to be greater in men, this association does not necessarily confirm causation. Based on our present results, we propose that healthy women have a platelet phenotype which is particularly sensitive to activation through thromboxane-mediated signaling which is blocked when aspirin irreversibly acetylates cyclooxygenase, and so healthy women may therefore derive more benefit from aspirin for primary prevention (29). When platelet thromboxane production is blocked, there is a reduction in thromboxane secretion and, platelet ‘auto-activation’, which is an important amplification step in thrombus formation, is attenuated (30). A very recent study in a murine mode of arterial thrombosis confirmed that lower doses of aspirin (81-100 mg) satisfactorily attenuate the thrombotic potential of platelets through thromboxane inhibition, while higher doses of aspirin, by preventing platelet prostacyclin production, promote bleeding (31). It is noteworthy, however, that the investigators only studied male mice. We therefore caution that there is neither clear evidence to suggest empirically increasing the dose of aspirin in men confers additional protective properties from thrombosis nor is there evidence to suggest that high dose aspirin promotes more bleeding in women. Conducting a study in healthy women and men with low (81 mg), intermediate (162 mg), and high (325 mg) dose aspirin, assessing thromboxane B_2_ to assess adherence and efficacy, then evaluating several methods of platelet activation through the thromboxane receptor will be required to truly prove whether a divergent platelet phenotype in women and men is responsible for the differences in cardiovascular outcomes observed with aspirin use.

We observed that platelet GPIIb/IIIa activation in healthy human platelets is greater in females. The CRUSADE study revealed female patients with NSTEMI treated with an intravenous GPIIb/IIIa antagonist have more bleeding than males, suggesting a sex-specific difference in GPIIb/IIIa signaling(17). At this time, intravenous GPIIb/IIIa antagonist dosing regimens do not account for gender-specific differences in platelet activation. Future investigations should focus on a gender-dependent effect of GPIIb/IIIa antagonists in the post-MI setting and on cardiovascular outcomes.

Off-target effects of anti-platelet agents and unexpected lack of efficacy in patients with certain demographic profiles or in certain clinical scenarios has been extensively reported. Single nucleotide polymorphisms in genes encoding enzymes for P2Y_12_ antagonist metabolism, opiate-mediated alterations in P2Y_12_ antagonist absorption, and metabolic disease-mediated alterations in the P2Y_12_ receptor conformation are proposed mechanism for failure of clopidogrel and ticagrelor to exert their anti-platelet affect (4,32,33). Enteric-coated aspirin resulting in delayed absorption, changes in the activity of drug transporters, and possible under-dosing are proposed mechanisms to explain an alteration in the efficacy of aspirin (19) (34). Determination of a gender-specific difference in these clinical situations as we present here is a logical extension of those previous studies. We previously reported that proteomic profiles, agonist sensitivity, and anti-platelet medication efficacy are altered in humans and in murine models of MI and peripheral artery disease (1,3). We now suggest, based on the present data, that platelets in patients with acute MI are primed toward augmented PAR signaling in men. Reasons for augmented platelet activation in males at the time of MI are unlikely to be explained only by sex-specific changes in platelet surface receptor density which we report here, but likely involve several alterations in post-receptor signal transduction pathways in male and female platelets. Further studies should focus on examining downstream (post-receptor) signal transduction pathways in platelets.

The present data supports our prior observations in a small patient cohort that platelets from patients with NSTEMI preferentially activate through PAR1 compared to platelets from patients with STEMI, though clinical and demographic variables to account for the observation were unclear (1). We now identify sex as an independent predictor of platelet PAR1 signaling following NSTEMI. The Acute Coronary Syndromes (TRACER) study suggested that a PAR1 antagonist added to dual anti-platelet therapy (DAPT) had some benefit in reducing adverse cardiovascular events with an increased risk of major bleeding (35), while a PAR1 antagonist alone in patients with established vascular disease significantly prevented arterial thrombosis and limb ischemia (36). Similarly, the MANAGE study showed that direct thrombin inhibition with dabigatran in patients with vascular disease and evidence of myocardial injury at the time of non-cardiac surgery conferred protection from thrombosis and vascular death (37). Focusing more on inhibiting platelet Protease Activated Receptors (PARs) or circulating thrombin as a mechanistic intervention—particularly in males in the peri-MI environment—is logical based on the current findings and previous investigations.

Using a Factor Xa inhibitor alone in the ATLAS ACS2 study (38) or in combination with aspirin in the COMPASS study, there was a dramatic reduction in atheroembolic events and improved mortality in patients with known vascular disease (39). It is noteworthy that these studies showed outcomes that were several-fold greater in men than women. The present study and our clinical experience is that twice as many males as females present with acute MI. Together, these observations raise the possibility that indirectly blocking thrombin with a Xa inhibitor may show the most survival benefit in males as reported (24,39). Moreover, since Xa inhibitors indirectly inhibit circulating downstream Factor IIa (thrombin) activity (40), it is striking that our present study confirmed in both humans and in mice that thrombin-mediated platelet activation is especially important in males in the immediate post-MI environment.

### STUDY LIMITATIONS

This investigation provides mechanistic insight that platelets from males and females respond differently to surface receptors for which drugs are available in health and following MI. A prospective investigation in which platelet thromboxane receptor signaling is blocked directly (aspirin) and platelet thrombin signaling is blocked directly (thrombin antagonist, PAR1 antagonist) or indirectly (Factor Xa inhibitor) will be required to assess whether sex-specific antithrombotic therapy should be employed acutely to affect mortality in patients with acute MI. Sex-specific differences in platelet signaling should also be examined in patients with chronic vascular disease such as CAD and PAD in whom recurrent atheroembolic events involving platelets are common.

### CONCLUSIONS

Inhibiting both platelet thrombin and thromboxane signaling in multiple vascular beds could be one mechanism for improving outcomes in stable vascular disease and in patients transitioning off dual anti-platelet therapy between 6 to 24 months post-MI. Our results indicate that greater attention should be directed towards whether changes in the platelet phenotype in women and men are responsible for differences in outcomes and responsiveness to anti-platelet medications.

### PERSPECTIVES

#### COMPETENCY IN MEDICAL KNOWLEDGE

Platelet thromboxane receptor and glycoprotein IIb/IIIa signaling were especially active in platelets from healthy females compared to males. Platelet PAR1 was strongly activated in males and attenuated in females at the time of MI compared to healthy conditions.

#### TRANSLATRIONAL OUTLOOK

Platelet phenotype and function are different in health and in the immediate post-MI environment for women and men, and therefore may shows differences in the response to anti-platelet agents. The predilection of platelets from healthy women to be activated by thromboxane pathway signaling may explain why aspirin appears to be more protective for stroke prevention compared to men. NSTEMI independently predicted platelet PAR1 as well as platelet thromboxane signaling but not P2Y_12_ receptor signaling, providing mechanistic as to why Factor Xa inhibitors in conjunction with aspirin are so effective in preventing recurrent atheroembolic events.

## Acknowledgements

We would like to Dr. Johns Gassler for advice on timing of blood collection, and Ms. Rachel Schmidt for technical assistance.

---

**Figure. S1:**
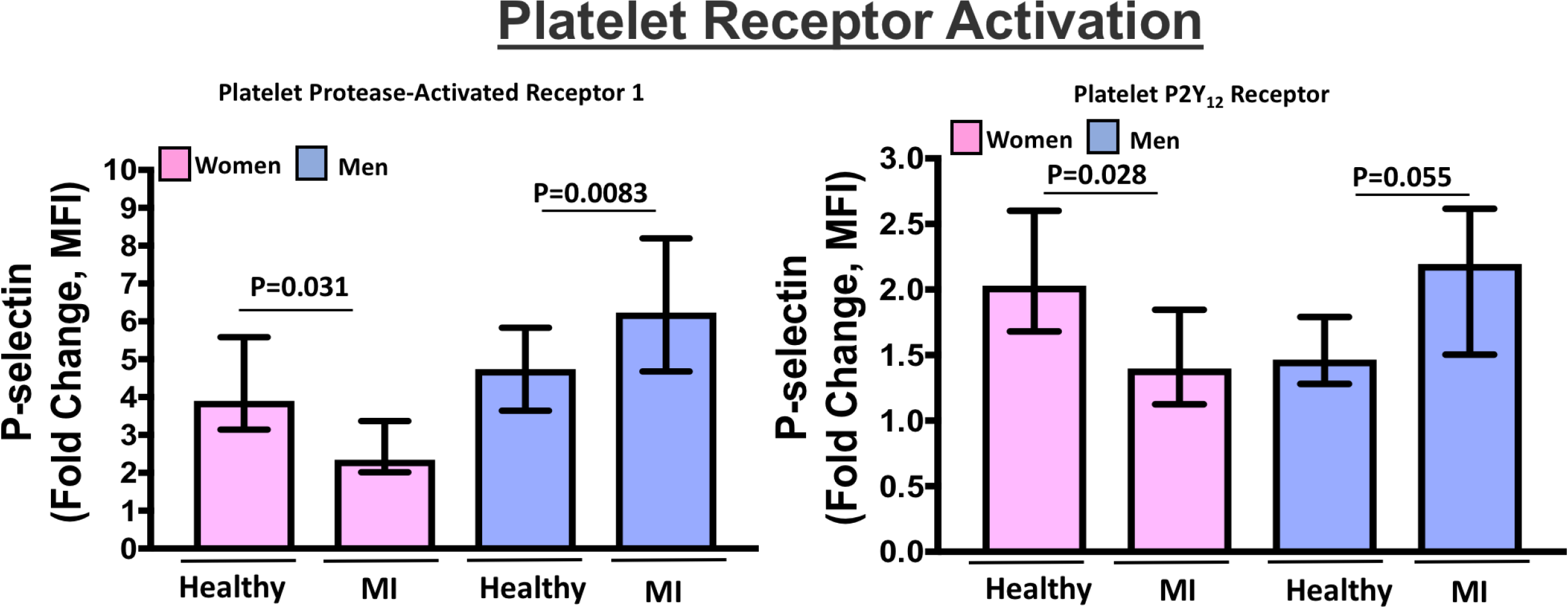
Agonist-mediated platelet activation in health and after Ml. Blood was drawn from healthy women and men and compared to blood drawn from patients with Ml (STEMI + NSTEMI) as soon as they were diagnosed (prior to coronary angiography, and prior to loading with a P2Y _12_ receptor antagonist). Platelet rich plasma was isolated and platelets were stimulated for 15 mins with an agonist for platelet PARl (TRAP-6, 10 µM) or the P2Y_12_ receptor (ADP, 10 µM). Platelets were labeled with a tagged antibody for p-selectin, and then analyzed by flow cytometry. Platelet function is represented as mean fold change in surface P-selectin by median fluorescence intensity (MFI) from baseline± 95% C.I. Differences between women and men for each agonist was assessed by the M ann-Whitney U test. Level of significance is noted above the graph.

**Figure. S2:**
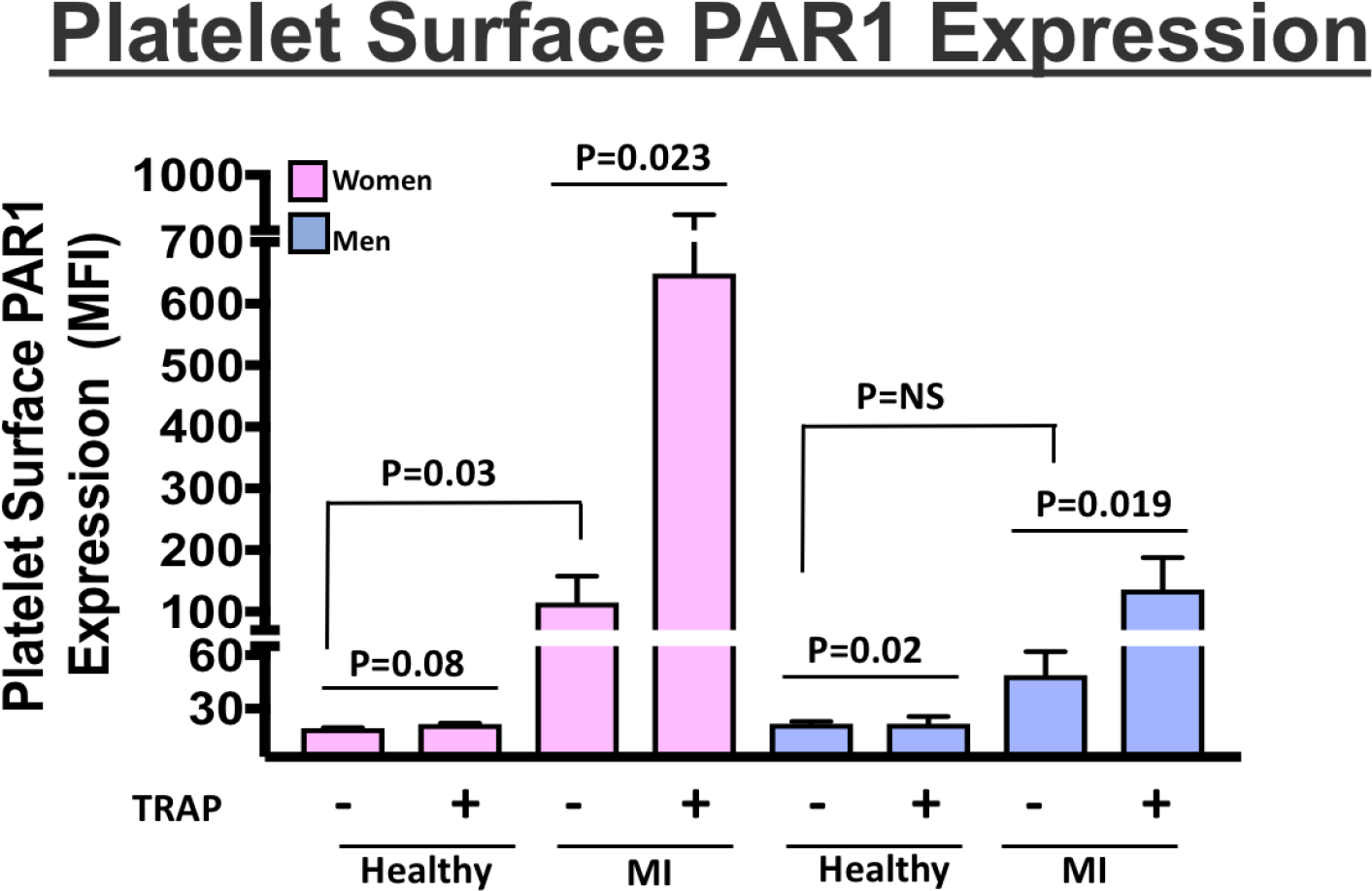
Platelet surface receptor expression after M.I. Blood was drawn patients as soon as they were diagnosed with NSTEMI, prior to coronary angiography, and prior to loading with a P2Y_12_ receptor antagonist. Platelet rich plasma was isolated and platelets were stimulated for 15 mins with an agonist for PARl (TRAP - 6, 10 µM). Platelets were labeled with a FITC-tagged antibody for PARl, and analyzed by flow cytometry. Platelet surface receptor density is represented as mean fluorescence intensity (MFI) ± SEM. Differences between groups was assessed by the Kruskal-Wallis test followed by Dunn ’s post test. Level of significance is noted above the graph.

**Figure. S3:**
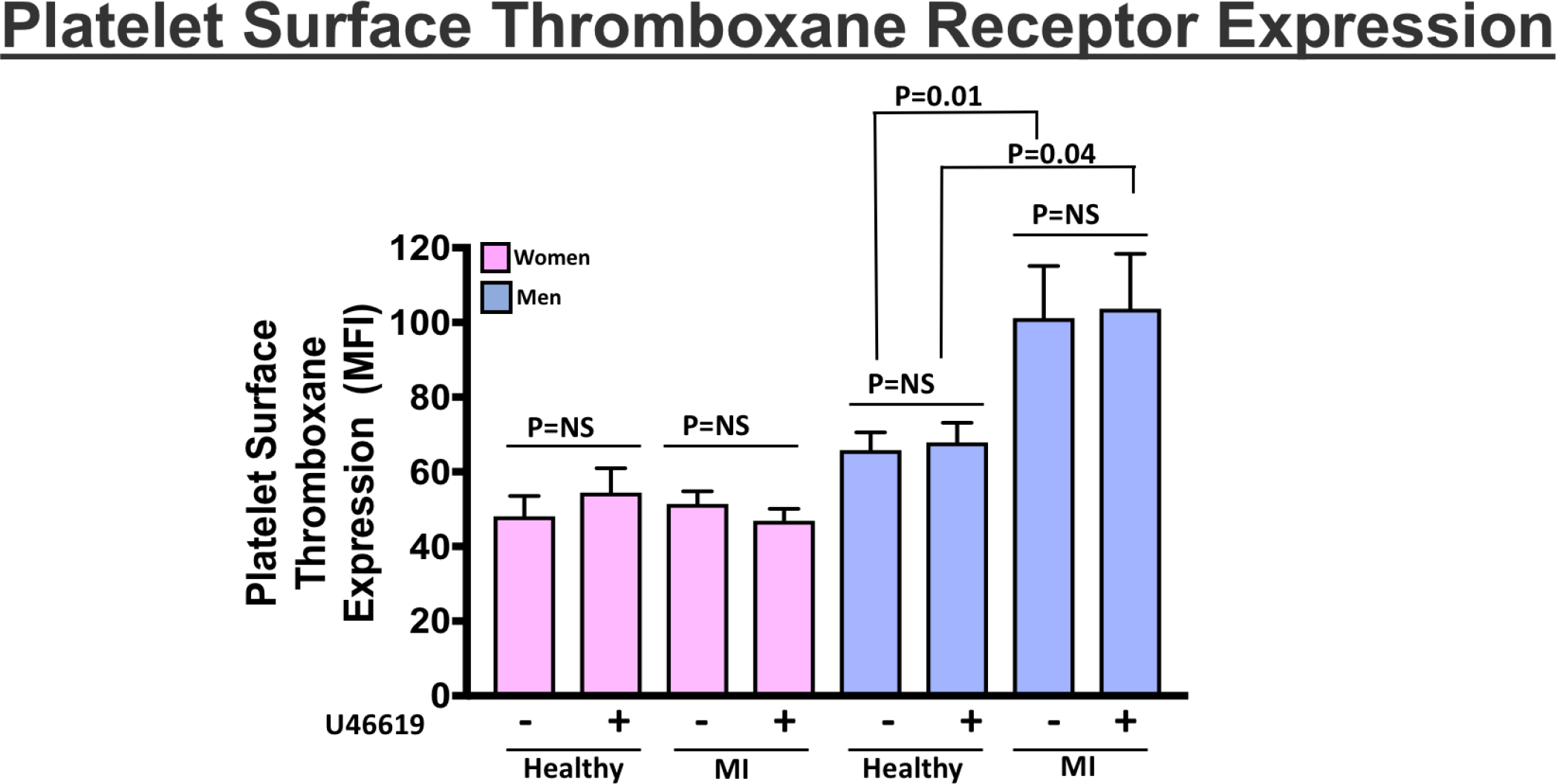
Platelet surface receptor expression after M.I. Blood was drawn patients as soon as they were diagnosed with NSTEMI, prior to coronary angiography, and prior to loading with a P2Y_12_ receptor antagonist. Platelet rich plasma was isolated and platelets were stimulated for 15 mins with an agonist for the thromboxane receptor (U46619, 10 µM). Platelets were labeled with a FITC-tagged antibody for thromboxane receptor, and analyzed by flow cytometry. Platelet surface receptor density is represented as mean fluorescence intensity (MFI) ± SEM. Differences between groups was assessed by the Kruskal­ Wallis test followed by Dunn’s post test. Level of significance is noted above the graph.

**Figure. S4:**
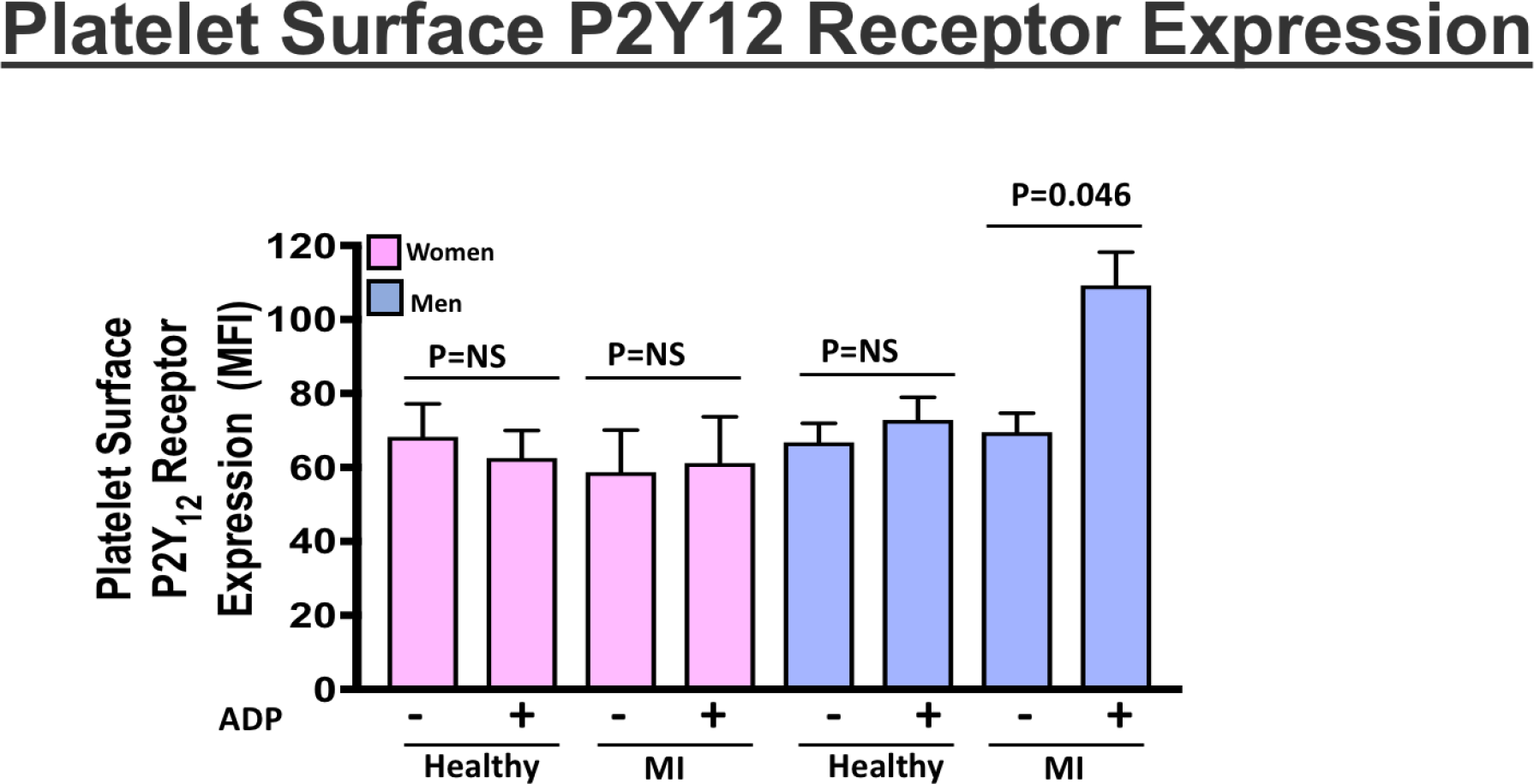
Platelet surface receptor expression after M.I. Blood was drawn patients as soon as they were diagnosed with NSTEMI, prior to coronary angiography, and prior to loading with a P2Y_12_ receptor antagonist. Platelet rich plasma was isolated and platelets were stimulated for 15 mins with an agonist for the thromboxane receptor (U46619, 10 µM). Platelets were labeled with a FITC-ta gged antibody for the P2Y_12_ receptor, and analyzed by flow cytometry. Platelet surface receptor density is represented as mean fluorescence intensity (MFI) ± SEM. Differences between groups was assessed by the Kruskal-Wallis test followed by Dunn’s post test. Level of significance is noted above the graph.

